# Antigen-Specific Immune Decoys Intercept and Exhaust Autoimmunity to Prevent Disease

**DOI:** 10.1101/662973

**Authors:** J. Daniel Griffin, Jimmy Y. Song, Aric Huang, Alexander R. Sedlacek, Kaitlin L. Flannagan, Cory J. Berkland

## Abstract

Relapsing-remitting patterns of many autoimmune diseases such as multiple sclerosis (MS) are perpetuated by a recurring circuit of adaptive immune cells that amplify in secondary lymphoid organs (SLOs) and traffic to compartments where antigen is abundant to elicit damage. Some of the most effective immunotherapies impede the migration of immune cells through this circuit, however, broadly suppressing immune cell migration can introduce life-threatening risks for patients. We developed antigen-specific immune decoys (ASIDs) to mimic tissues targeted in autoimmunity and selectively intercept autoimmune cells to preserve host tissue. Using Experimental Autoimmune Encephalomyelitis (EAE) as a model, we conjugated autoantigen PLP_139-151_ to a microporous collagen scaffold. By subcutaneously implanting ASIDs after induction but prior to the onset of symptoms, mice were protected from paralysis. ASID implants were rich with autoimmune cells, however, reactivity to cognate antigen was substantially diminished and apoptosis was prevalent. ASID-implanted mice consistently exhibited engorged spleens when disease normally peaked. In addition, splenocyte antigen-presenting cells were highly activated in response to PLP rechallenge, but CD3+ and CD19+ effector subsets were significantly decreased, suggesting exhaustion. ASID-implanted mice never developed EAE relapse symptoms even though the ASID material had long since degraded, suggesting exhausted autoimmune cells did not recover functionality. Together, data suggested ASIDs were able to sequester and exhaust immune cells in an antigen-specific fashion, thus offering a compelling approach to inhibit the migration circuit underlying autoimmunity.

## MAIN

Central to the propagation of adaptive immunity is the maturation of antigen-specific responses in secondary lymphoid organs (SLOs) such as the spleen and lymph nodes [1–3]. Following these initiatory events, antigen-specific populations are amplified, egress from SLOs, and traffic to antigen-rich locales [4, 5] for further stimulation and effector action [6–8]. Acute adaptive immunity is robust, but short-lived for the rapid clearance of pathogens and abnormal cells. Following the steep deployment of a response, immune cells can become exhausted as a checkpoint mechanism. Under these conditions, functional capacity is diminished and apoptosis may ensue [9–11]. Together, the migratory circuit between SLOs and diseased tissue coupled with the exhaustible nature of immunity is a hallmark of relapsing-remitting autoimmune diseases such as multiple sclerosis (MS) [12–15]. In such diseases and their experimental models, a vicious cycle is entrenched where aberrant autoantigen-specific effectors are primed, trafficked to disease-specific sites, and further stimulated to cause damage before returning to SLOs and regaining functionality [16–18].

Some of the most clinically effective immunotherapies for MS work by blockading the migratory circuit between peripheral SLOs and the central nervous system [19, 20]. Preventing egress from SLOs or blocking infiltration of myelin-specific immune populations can preserve host tissue integrity and stifle autoimmune destruction. These disease-modifying therapies are, however, nonspecific in their mechanism. The trafficking of entire populations of immune cells is broadly hindered, which severely handicaps both disease-relevant and healthy immune functions alike. Patients receiving such immunotherapies are subjected to risks of life-threatening infection that can far outweigh therapeutic benefits [21–23]. Antigen-specific immunotherapies may offer an appealing improvement to the state of conventional treatments, promising to selectively disarm autoreactive cells while preserving healthy immune function.

Conventional migration-inhibiting immunotherapies broadly block cellular egress from lymph nodes [19] or extravasation across the blood brain barrier [20]. Conversely, an antigen-specific approach may necessitate selectively attracting autoimmune cells or autoantibodies to a surrogate locale for sequestration. Engineered implantable biomaterials have emerged for evoking therapeutic, antigen-specific immune responses against cancer and autoimmunity alike [24–30]. These materials create a unique niche [31–36] of antigen to recruit and tune antigen-presenting populations for activation or tolerance, often by including a stimulatory adjuvant or immunosuppressive drug [37–45].

We designed and tested a novel biomaterial, which mimics etiologic autoantigens via covalent surface conjugation of a peptide epitope to microporous collagen sponges. Antige n-Specific Immune Decoys (ASIDs) were designed to short-circuit the SLO-host tissue circuit by sequestering autoreactive effector cells from circulation, ultimately preserving host organs. Using murine experimental autoimmune encephalomyelitis (EAE) as a model, we set forth to evaluate the prophylactic, subcutaneous implantation of ASIDs for preventing paralysis *in vivo*. Analyzing cell isolates from both sponges and spleens, we explored the hypothesis that these constructs could work by intercepting T cell proliferation in the periphery. Ultimately, we discovered that ASIDs overstimulated immunity and exploited antigen-specific exhaustion as a mechanism for therapeutic effect (**Fig. 1**).

**Figure 1.**
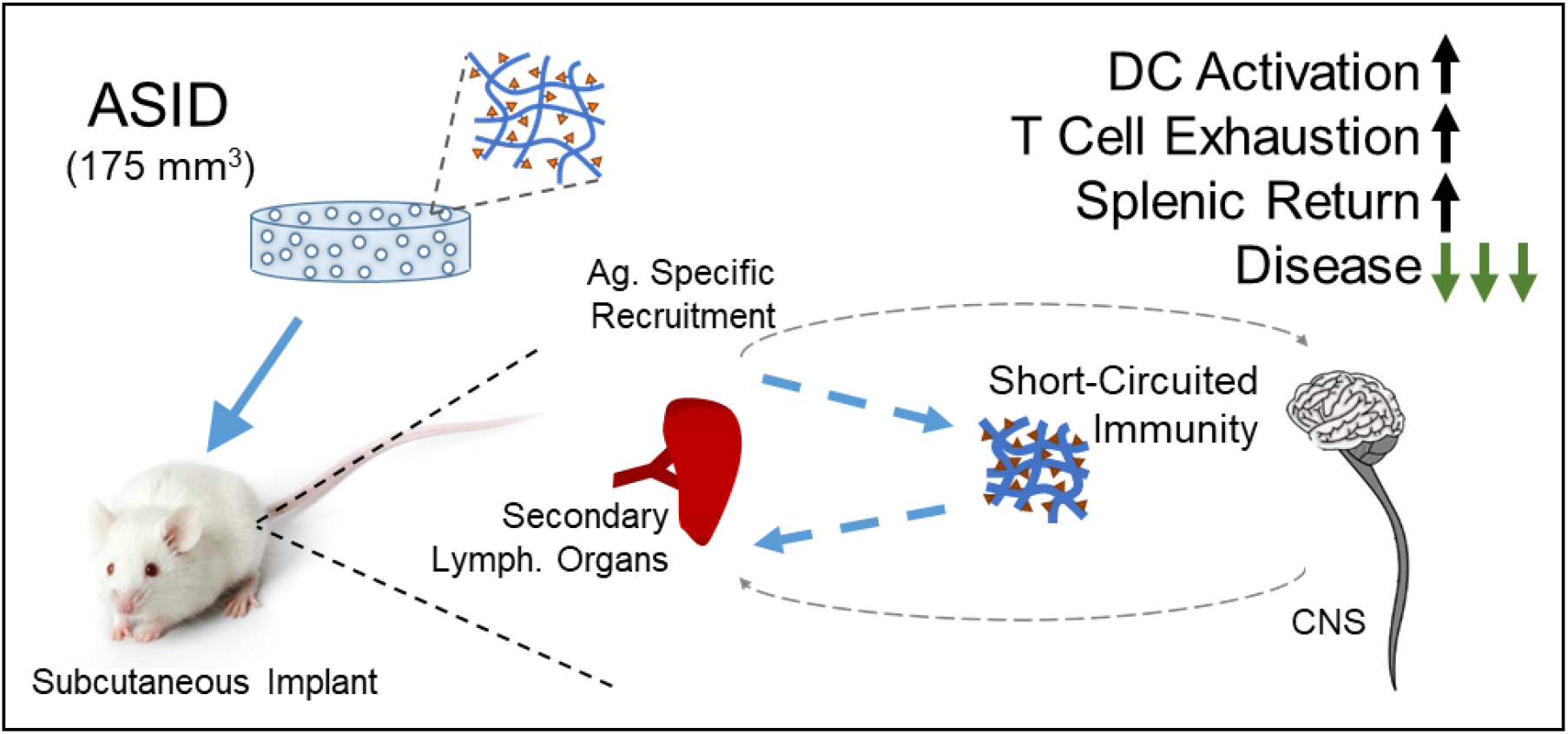
Autoimmunity is primed in secondary lymphoid organs and travels to disease-implicated tissues. ASIDs were designed to intercept this response and exhaust it prematurely, leaving host tissue intact.

## RESULTS

### Synthesis and Functional Characterization of ASIDs

We designed ASIDs as an implantable depot displaying covalently-coupled autoantigen. Microporous collagen sponges were selected as a scaffolding material due to immunologically inert nature and widespread clinical utility since the 1970s [46]. We hypothesized primary amine moieties would be chemically accessible on the surface of these collagenous materials (**Fig. 2a**) and pore diameters of ~200 μm would facilitate unhindered cell infiltration and nutrient transfer (**Fig. 2b**). To qualitatively assess surface accessibility of primary amines, a bifunctional NHS-PEG4-Azide linker was installed to collagen sponges, and rhodamine-alkyne was successfully conjugated using copper (I)-catalyzed azide-alkyne cycloaddition (**Fig. 2c**). Likewise using the same chemistry, a homopropargyl version of model autoantigen PLP_139-151_ (hpPLP) was coupled to the collagen sponges to synthesize ASIDs. PLP conjugation was confirmed via RP-HPLC.

**Figure 2.**
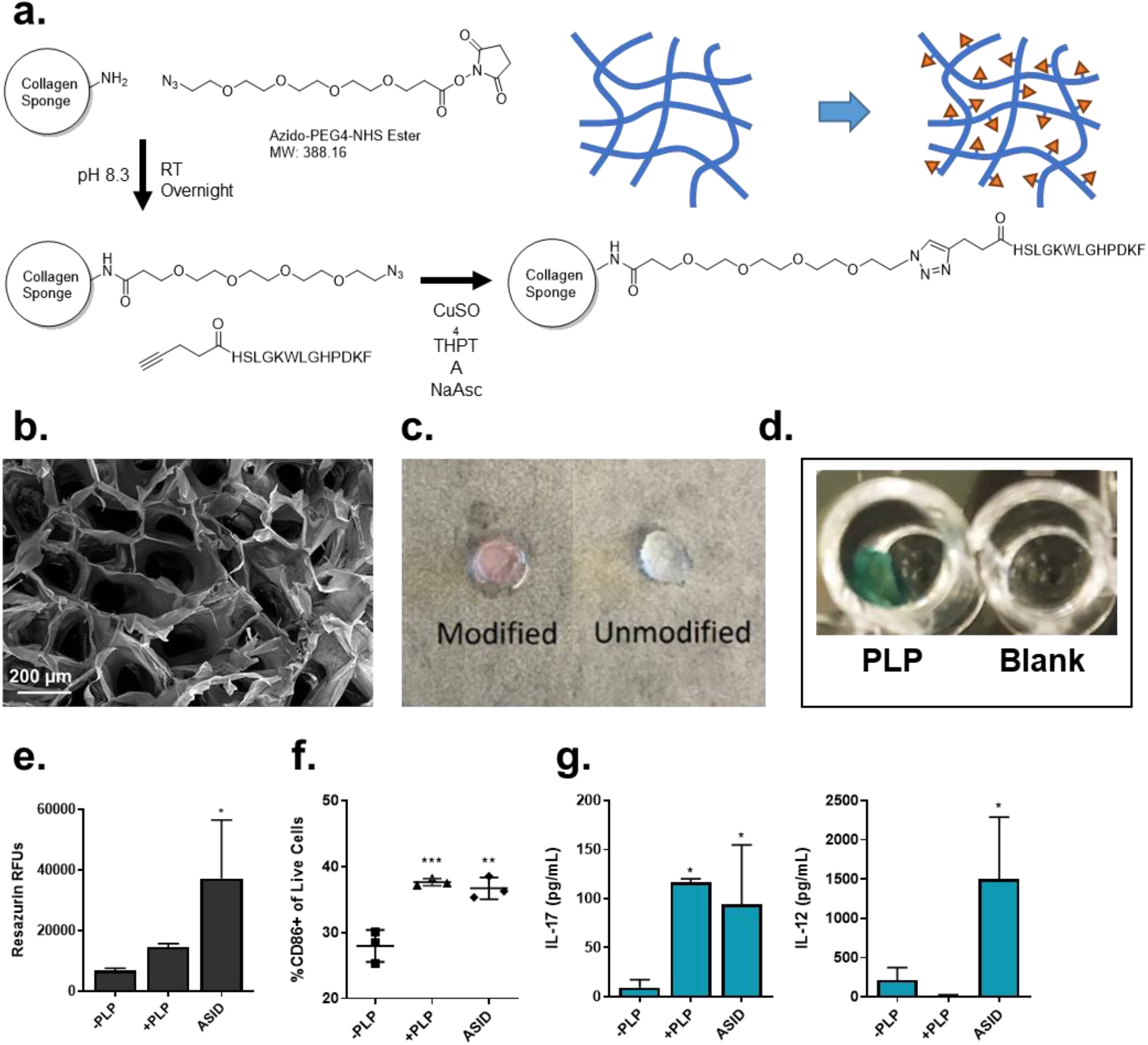
Synthesis and characterization of immunologically active PLP-conjugated ASIDs. **a)** The synthesis reaction scheme modified surface primary amines on collagen sponges with an azide-functional PEG linker subsequently clicked to homopropargyl PLP_139-151_. **b)** Microporous collagen sponges were used with an average pore size of approximately 200 μm. **c)** To visualize surface accessibility of primary amine moieties, rhodamine-alkyne was installed (left) and imaged compared to a Blank sponge (right), where dye conjugation could be confirmed. **d)** ASID with surface-conjugated PLP (left) captured anti-PLP IgG (note blue color) from mouse serum and a Blank collagen sponge (right) did not. **e)** Day 12 EAE Splenocyte metabolism was measured after incubation with vehicle (-PLP), soluble antigen (+PLP), or ASID. ASIDs significantly elevated metabolism over vehicle-treated splenocytes after 96 hours. **f)** Under the same conditions, CD86 expression was measured by flow cytometry, where ASIDs likewise upregulated costimulation in a manner similar to that of soluble antigen. **g)** In the same experiment, quantification of cytokine production showed that inflammatory IL-17 (left) was elevated by both +PLP and ASID treatment, however APC-indicating IL-12 (right) was only stimulated by incubation with ASIDs. (Statistical analysis was performed against –PLP as a control. n = 3/group, *p < 0.05, **p < 0.01, ***p < 0.001).

The functionality of conjugated autoantigen was then validated. ASID and Blank collagen sponges were each incubated in anti-PLP IgG-positive mouse serum. HRP-conjugated anti-mouse IgG was used to detect bound autoantibody (**Fig. 2d**), and the ASID, but not Blank sponge, showed successful capture. Additionally, EAE splenocytes were harvested at peak of disease and cultured *ex vivo* with ASIDs. Functional outcomes were compared with cells stimulated with 25 μM soluble PLP_139-151_ as well as vehicle control (**Fig. 2e-g**). After 96 hours, splenocytes cultured with ASID exhibited a significantly elevated metabolism over vehicle control (**Fig. 2e**), and CD86 expression levels were analogous to cells stimulated with soluble antigen (**Fig. 2f**). Likewise, soluble PLP and ASIDs generated robust IL-17 secretion, though only ASID culturing enabled high IL-12 after the incubation, indicating sustained APC activity (**Fig. 2g**).

### Subcutaneous Implantation of ASIDs prevents EAE in vivo

To investigate ASID therapeutic capacity, we subcutaneously implanted EAE mice with these constructs between the shoulder blades on day 7 post-induction, prior to the onset of symptoms (**Fig. 3a**). Healthy mice were also implanted with ASID to assess whether decoys themselves would stimulate an immune response. All implanted sponge constructs were sterilized and soaked overnight in a solution of 1200 ng/mL mouse GM-CSF prior to implantation to create impetus for cell infiltration and to normalize for inflammatory discrepancies that may have arisen from variability in surgery. ASIDs highly suppressed EAE *in vivo*, both in terms of significant scoring reduction as well as weight loss (**Fig. 3b-d**). Only two of four ASID mice showed any EAE symptoms at all, and none exhibited substantial paralysis which is defined as a clinical score of 2 or greater (**Fig. 3e**). Blank sponge implantation and Mock Surgery had no significant clinical effect on disease course, as these mice did not vary as compared to EAE alone with no surgical procedure or treatment. Control disease in this study was severe, as one mouse from the Blank group and one mouse from the Mock Surgery group each succumbed to disease at its peak. Strikingly, weight gain in EAE mice treated with ASID mirrored that of the healthy mice that were completely unhindered by disease (**Fig. 3c**).

**Figure 3.**
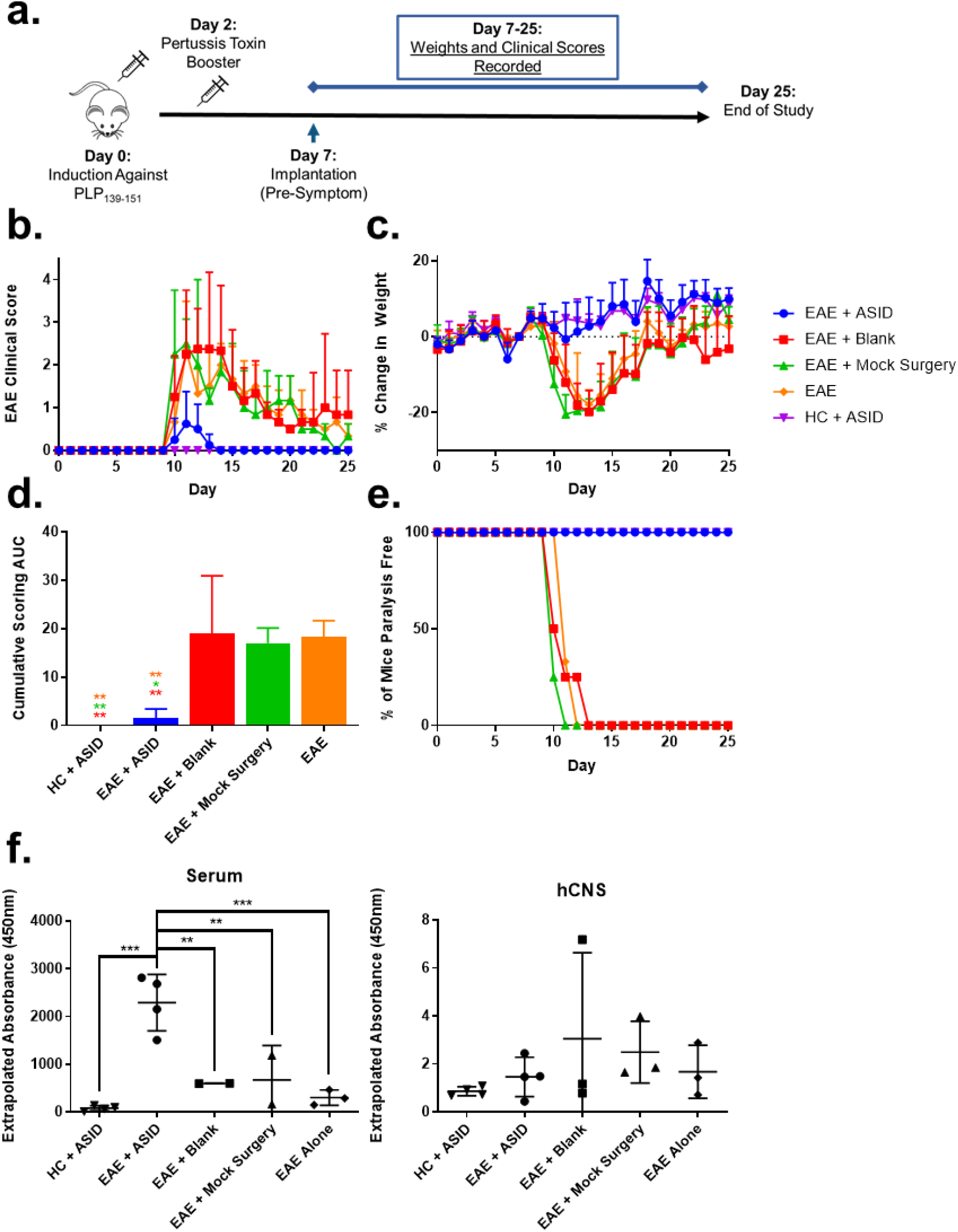
Therapeutic Evaluation of ASIDs against EAE in vivo. **a)** Mice were induced with EAE (excluding the HC, healthy controls group). On day 7, before symptoms began, mice were subcutaneously implanted with either an ASID or Blank Collagen Sponge, given Mock Surgery, or no treatment (EAE alone). **b)** Clinical Scoring and **c)** Weight change data were collected over the course of 25 days and demonstrated that ASID implantation significantly suppressed disease and did not induce autoimmunity in healthy mice. **d)** Cumulative clinical score was calculated and compared between groups, reinforcing the significance of ASID effect. **e)** Additionally, disease incidence was stratified by presence of disease at or above a clinical score of 2, where ASID implantation completely prevented incidence of severe symptoms. **f)** After the study, serum and hCNS were collected from mice, and anti-PLP IgG titer was compared. ASID-implanted EAE mice exhibited a significantly elevated serum titer of anti-PLP IgG, but there was no change in hCNS titer (n = 3-4/group, *p < 0.05, **p < 0.01, ***p < 0.001).

At the conclusion of the study, serum and whole brain homogenates were collected. Anti-PLP_139-151_ IgG ELISA was used as previously described [47–49] to detect relative autoantibody titers between the periphery (serum) and CNS (homogenized whole brain, **Fig. 3f, Supp. Fig. 1**). EAE mice implanted with an ASID presented with an elevated serum anti-PLP titer that did not translate to significant increases in CNS autoantibody.

### ASID Infiltrates undergo Apoptosis upon PLP Rechallenge

We hypothesized the persistent, antigen-specific depot in ASIDs could selectively sequester autoreactive cell populations responsible for propagating autoimmunity to prevent disease. To evaluate this proposed decoy mechanism, we implanted EAE mice with *both* a Blank collagen sponge as well as an ASID on day 7 post-induction **(Fig. 4a**). Sponges and spleens were harvested at typical disease onset (day 10) or peak of disease (day 14). Cells were isolated and rechallenged with 25 μM PLP for 96 hours.

**Figure 4.**
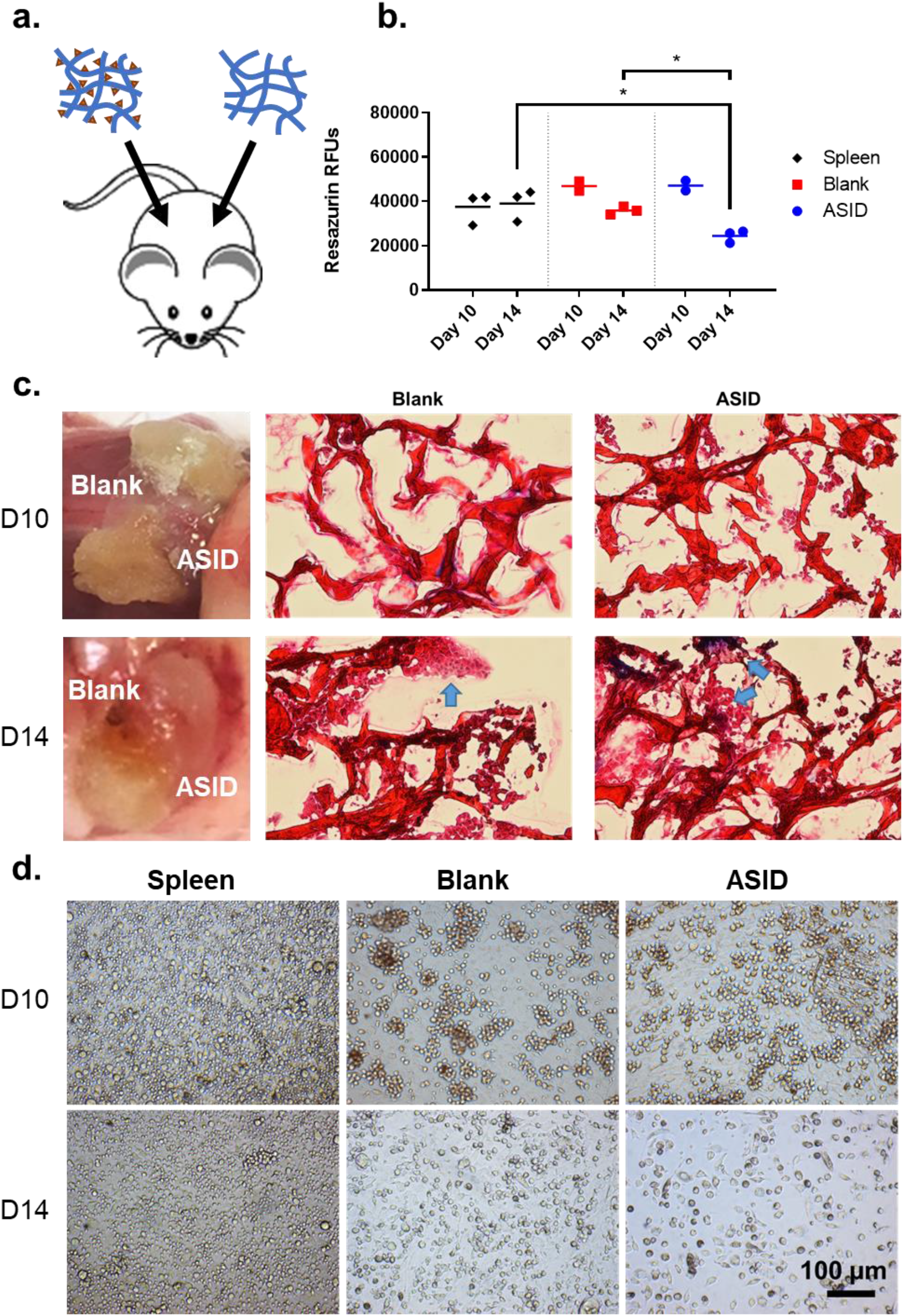
Same-animal sponge infiltrate analysis. **a)** On day 7 post-induction, EAE mice received both an ASID and Blank Sponge separately placed in the subcutaneous space. **b)** At both onset (day 10) and peak (day 14) of disease, sponges and spleens were retrieved and challenged with 25 μM PLP. After 96 hours, metabolism was measured, and showed that while few differences were appreciable at day 10, the day 14 timepoint demonstrated that metabolism was significantly decreased in ASID cell infiltrates. **c)** Resected sponges were imaged upon retrieval and cryosectioned for trichrome histology on day 10 (top) and day 14 (bottom). Here, healthy and ordered cell structures were observed in the Blank construct by day 14, but ASID infiltrates were disordered and misshapen. **d)** These observations were conserved in splenocytes (left) and spongeocytes (Blank, middle, and ASID, right) after being isolated from the same mouse and rechallenged with 25 μM PLP. After 96 hours, splenocytes and Blank sponge infiltrates appeared largely healthy, however death was abundantly prevalent in ASID isolates (n = 3 biological replicates per group, *p < 0.05).

At day 10, there were no significant differences in resazurin cell metabolism across splenocytes and sponge isolates (**Fig. 4b**), though ASIDs were considerably more degraded than Blank sponges (**Fig. 4c**). At day 14, metabolism in ASID infiltrates was roughly halved as compared to the Blank sponge and spleen **(Fig. 4b**). These results were initially surprising as we expected PLP-specific cells isolated from ASIDs to elicit a greater metabolic response upon rechallenge. To further investigate, harvested sponges were cryosectioned and Masson’s trichrome staining was used to assess morphological differences (**Fig. 4c**). Reflecting resazurin metabolism data, few differe nces were observable at day 10 (though ASID sections showed relatively more cell infiltrates). In contrast, ASID infiltrates appeared disjointed and unhealthy at the day 14 timepoint, while Blank sponge sections showed apparently healthy, ordered cell structures. Bright field microscopy of isolated splenocytes and spongeocytes after 96-hour PLP rechallenge corroborated apoptotic allusions. ASID isolates at the day 14 timepoint were clearly dying off in comparison to the Blank isolates and splenocytes.

### ASIDs Return Exhausted Immune Cells to Secondary Lymphoid Organs

In the same-anima l implant experiment (**Fig. 4**), we noticed that secondary lymphoid organs appeared substantially engorged at times in disease course when splenopenia is typical, due to autoimmune cell egress to the CNS (**Supp. Fig. 2**). To further probe this phenomenon as a potential mechanistic underpinning of ASID efficacy, we implanted EAE mice with either an ASID, Blank sponge, or Mock Surgery on day 7 post-induction and harvested at peak of disease on day 14 (**Fig. 5a**). ASID-implanted mouse spleens were larger than those of the other groups by a factor of 4, and a similar size discrepancy in lymph nodes was evident (**Fig. 5b, Supp. Fig. 3**). The largest numbers of splenocytes were also harvested from these spleens, where yields averaged a multiple of two and four times those from Mock and Blank controls, respectively. Cells were also isolated from sponges, and ASID isolates numbered roughly 5 million cells per construct while Blank infiltrates were on the order of 10,000 cells per sponge (**Fig. 5b**).

**Figure 5.**
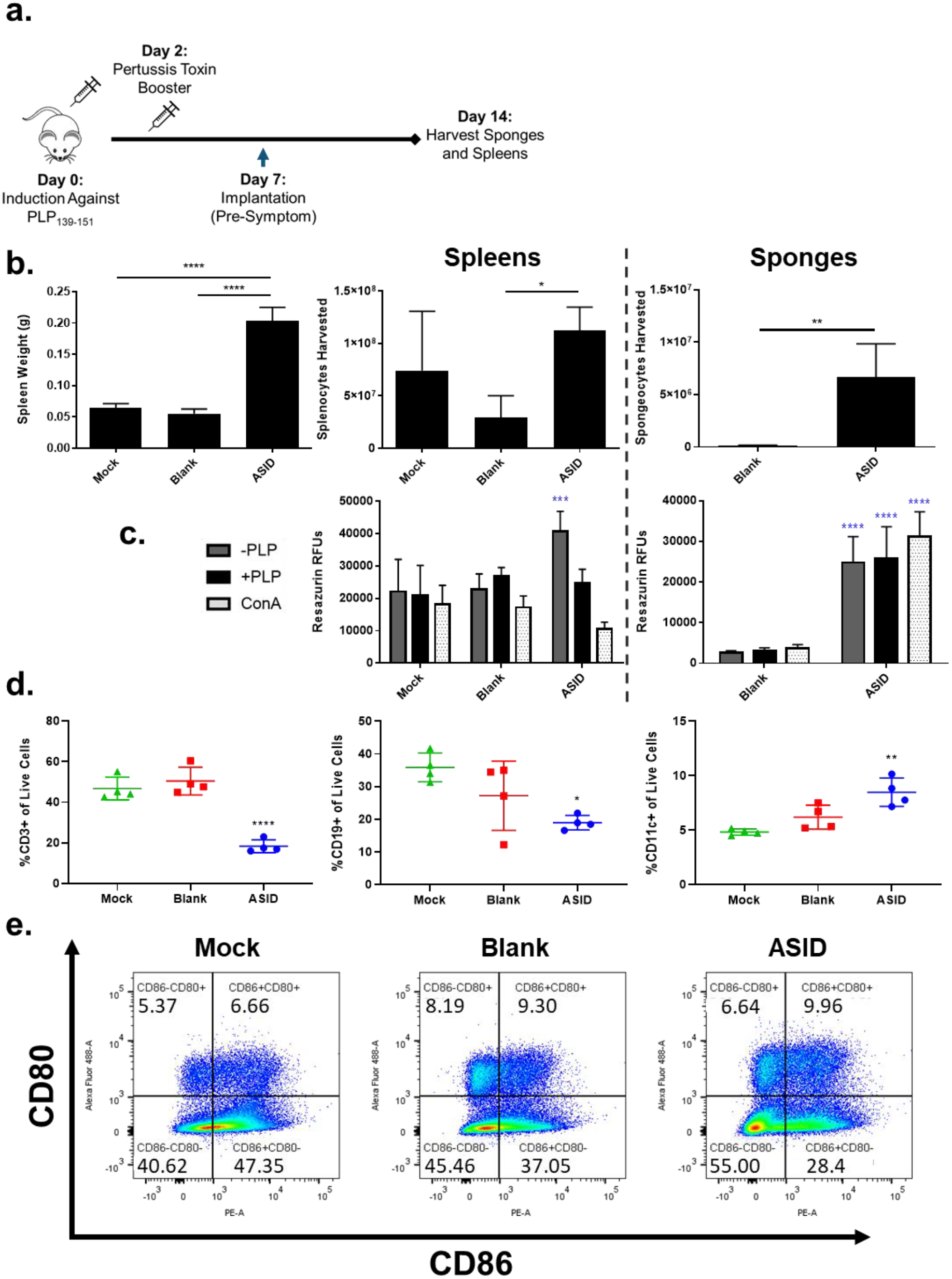
Differences in composition at peak of disease. **a)** EAE mice were implanted with ASID, Blank Sponge, or Mock Surgery. On day 14, spleens were harvested. **b)** Spleen weights were compared at harvest (left) in addition to isolated splenocyte (middle) and spongeocyte (right) counts. ASIDs both recruited the most spongeocytes and induced significant spleen engorgement. **c)** Cell isolates from spleens (left) and sponges (right) were incubated with vehicle (-PLP), 25 μM PLP (+PLP), or 2.5 μg/mL Concanavalin A (ConA) for 96 hours and cell metabolism was measured. ASID splenocytes exhibited high metabolism when incubated with vehicle, but PLP rechallenge diminished this elevation. In spongeocytes, metabolism was largely conserved across all three treatments. **d)** Flow cytometry was used to phenotype splenocytes for CD3 (left), CD19 (middle), and CD11c (right), and splenocytes from ASID implanted mice were found to be relatively depleted of T and B cell subsets. Additionally, CD11c+ populations were increased in ASID spleens as compared to controls. **e)** CD80 and CD86 were also observed between groups, where CD86+CD80+ costimulation was slightly elevated by Blank sponges and ASIDs. (Statistical analysis was performed against the Mock group as a control. n = 4 biological replicates group, *p < 0.05, **p < 0.01, ***p < 0.001, ****p<0.0001).

Isolated splenocytes and spongeocytes were rechallenged with media alone, 25 μM PLP, or 2.5 μg/mL of mitogenic Concanavalin A (ConA) for 96 hours (**Fig. 5c**). Unchallenged ASID splenocytes showed a significantly higher level of metabolism after the incubation (compared to controls), but the inclusion of PLP or ConA abrogated this increase. In the sponges, ASID isolates exhibited metabolism that was not exhaustive in response to PLP or ConA, and was significa ntly elevated over Blank sponge isolates, but these cells were much lower in number as mentioned above (**Fig. 5b**).

Flow cytometry was used to phenotype splenocytes immediately after harvest (**Fig. 5d, e)**. Moving from our previous same-animal experiment and observations of enlarged spleens, we hypothesized that ASIDs may have worked to evoke antigen overstimulation to exhaust the EAE autoimmune response and prematurely return perpetrating cells to secondary lympho id organs. In accordance with our hypothesis, we found T (CD3+) and B (CD19+) cell populations to be significantly depleted in ASID spleens compared to those of Mock Surgery (**Fig. 5d**). Seemingly conflicting with these decreased levels of effector subsets, though, was an increase in CD11c+ APCs (**Fig. 5d**) and comparable levels of CD86+CD80+ costimulatory expression (**Fig. 5e**).

### Antigen-Specific Exhaustion Persists after PLP rechallenge

Moving from baseline phenotyping, day 14 splenocytes were also rechallenged with 25 μM PLP for 96 hours. After the incubation, many baseline trends persisted (**Fig. 6a**). ASID splenocyte CD3+ T cells remained lower in proportion to controls both with and without rechallenge, though CD19+ B cells were relatively similar at this time point. CD11c+ APCs were significantly elevated when incubated with vehicle media, but not significantly different when challenged with antigen. However, an inverted antigen rechallenge trend was appreciable; the inclusion of PLP led to expanded CD11c+ populations in Mock and Blank controls, but led to a decrease when included with ASID splenocytes **(Fig. 6a**). Despite this decrease, the CD11c+ proportion of rechallenged ASID splenocytes was slightly higher than those of the controls. Costimulation was promoted upon rechallenge without exception across groups, but PLP rechallenge in ASID splenocytes promoted exorbitantly higher levels of CD80+ as well as CD86+CD80+ double positive cells (**Fig. 6a**).

**Figure 6.**
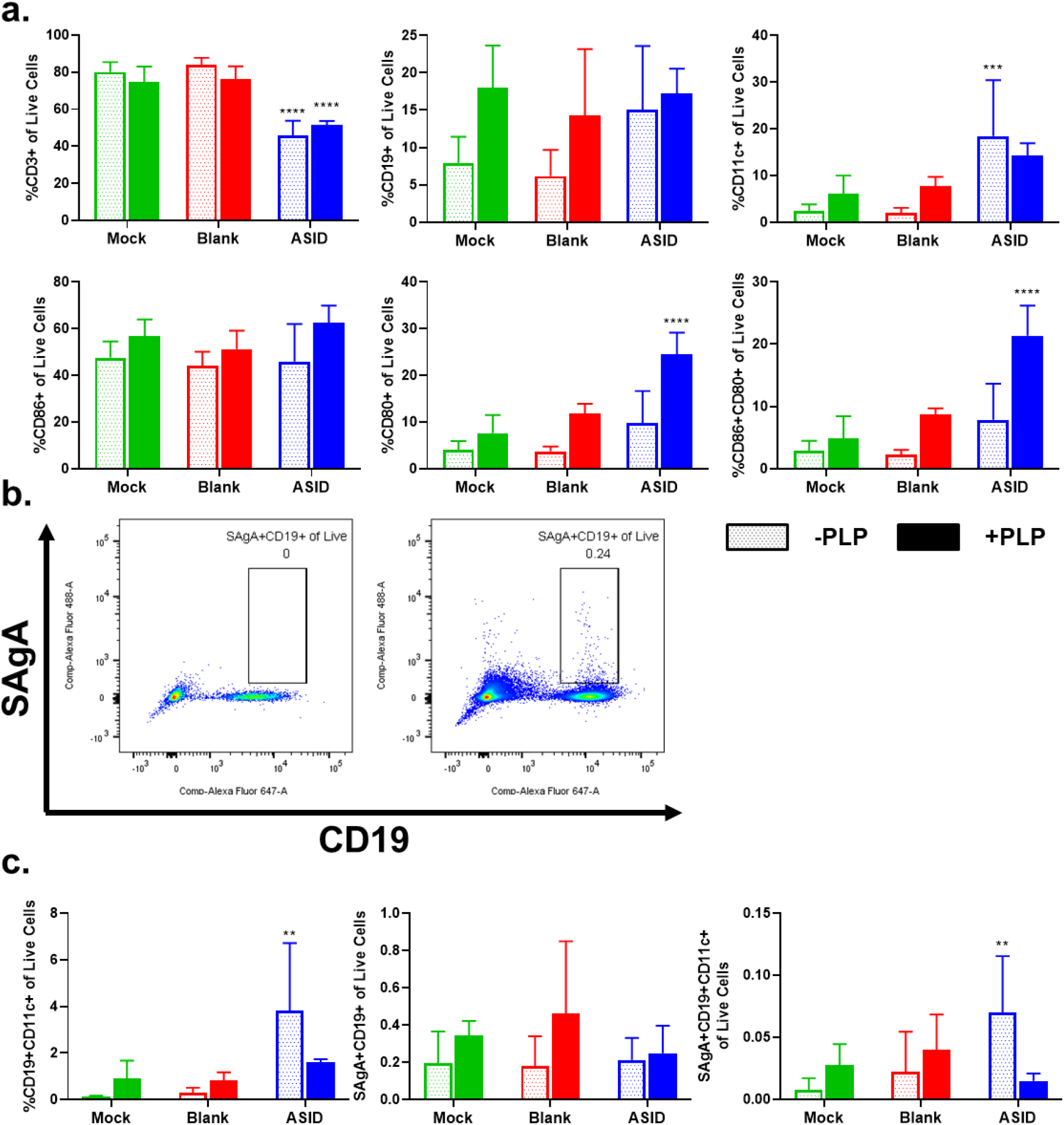
PLP rechallenge response among harvested splenocytes. **a)** From the previous experiment, excess splenocytes were phenotyped after 96 hours of vehicle (-PLP, open bars) or 25 μM PLP treatment (+PLP, solid bars) for CD3, CD19, and CD11c (top row, left to right) as well as costimulatory markers CD86, CD80 and double positive CD86+CD80+ populations (bottom row, left to right). Similar trends to the baseline analysis were appreciable; CD3+ T cell proportions were suppressed in ASID spleens, but CD11c+ populations were slightly more prevalent. **b)** SAgAs were used to reliably tag antigen-specific B-cells and ultimately **c)** probe disease-relevant ABCs (left), SAgA+ B cells (middle), and SAgA+ ABCs (right). This analysis provided insight to show that within ASID splenocytes, the ABC subset showed a considerable decrease in prevalence in response to antigen rechallenge. (Statistical analysis was performed against the Mock group as a control. n = 4 biological replicates per group, *p < 0.05, **p < 0.01, ***p < 0.001, ****p<0.0001).

One major limitation of the EAE model is a lack of tools for probing antigen-specificity. In recent years, our group has published a multivalent nanomaterial known as the soluble antigen array (SAgA), which has been developed as an antigen-specific immunotherapy [50–53]. SAgAs evoke efficacy by specifically engaging surface-bound B cell receptors [54–57]. In this study, we hypothesized that fluorescently labeled SAgAs could identify antigen-specific B cells much in the same way that MHC-tetramers are employed for T cells. By including SAgAs with other fluorescent antibodies, we developed a method in which antigen-specific cells could be identified by gating on CD19 (**Fig. 6b, Supp. Fig 4**). Extrapolating this protocol allowed identification of antigen-specific subsets of B cells in EAE splenocytes **(Fig. 6c**). CD19+CD11c+ autoimmune-associated B cells (ABCs) have been reported as potent, antigen-specific instigators of immunity in EAE and autoimmune disease alike [57–59]. Interestingly, ASID splenocytes showed significantly elevated proportions of this population when incubated with vehicle (**Fig. 6c**), though much like resazurin (**Fig. 4b, Fig. 5c**), these cells were diminished by PLP rechallenge. The SAgA probe verified this trend was conserved within the antigen-specific CD19+CD11c+ ABC subset, but not the broader CD19+ B cell population.

### Pro-inflammatory Cytokines are Elevated in ASID-Treated Splenocytes at Day 14 and 25

In addition to phenotypic changes in response to antigen rechallenge, we assessed cytokine responses in splenocytes harvested on day 14 or 25 and cultured with 25 μM PLP or vehicle for 96 hours (+PLP reported in **Fig. 7a,**-PLP and absolute concentrations in **Supp. Fig. 7**). At day 14, ASID mice showed increased in pro-inflammatory cytokines GM-CSF, IFN-γ, and IL-6 as compared to the Mock Surgery group, while splenocytes from mice implanted with a Blank sponge largely secreted less of each analyte screened. At day 25, ASID-treated splenocytes still exhibited elevated levels of pro-inflammatory cytokines IFN-γ, IL-15, IL-17, and IL-6, though at this point, anti-inflammatory IL-10 was most profoundly upregulated.

**Figure 7.**
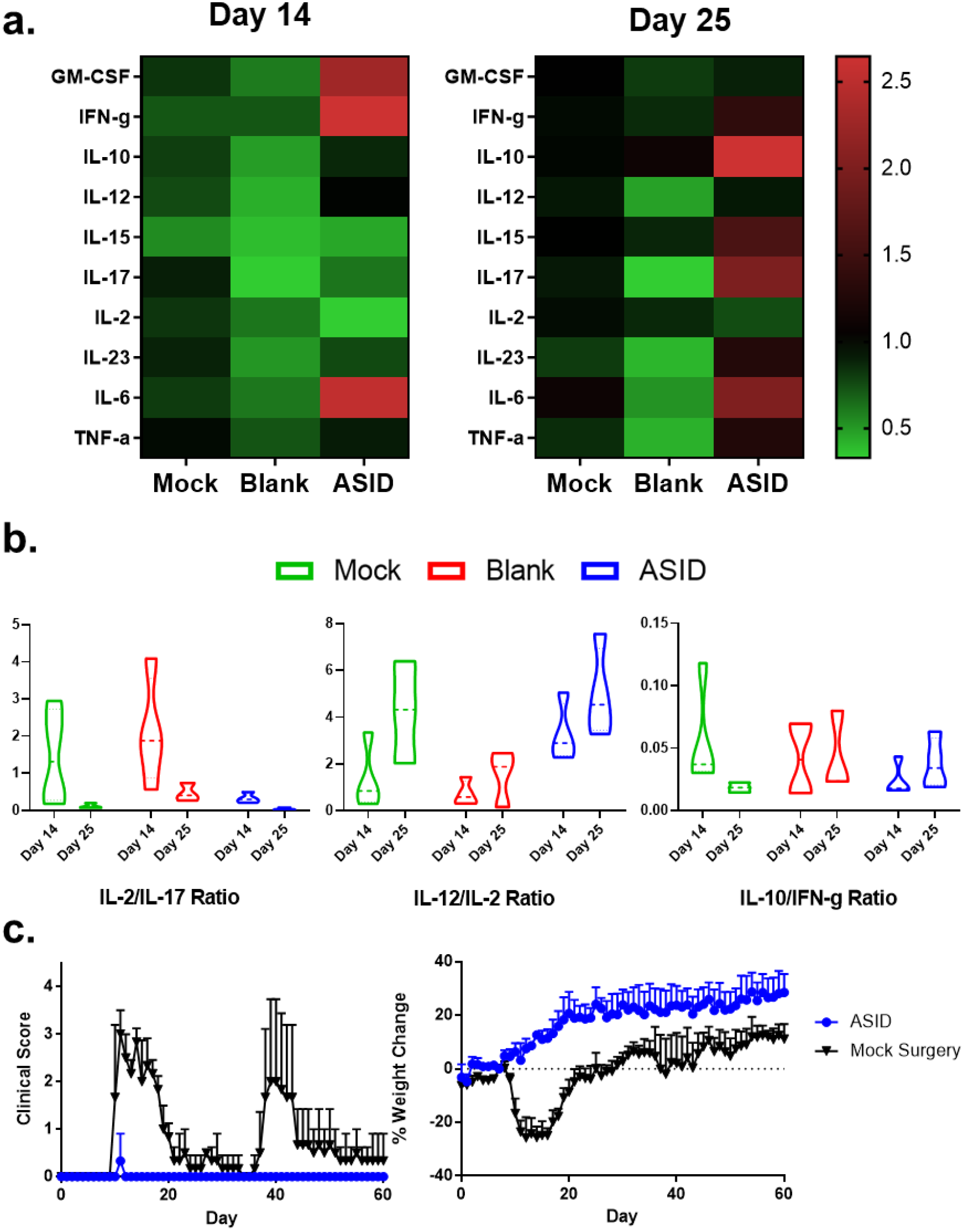
Cytokinechanges in PLPchallenged splenocytes. **a)** Day 14 (left) and day 25 splenocytes (right) were challenged for 96 hours with 25 μM PLP, and changes are expressed in terms of median fold-change with respect to Mock as a control. At day 14, inflammatory cytokines GM-CSF, IFN-γ, and IL-6 abounded in ASID splenocytes, though IL-2 was suppressed. At day 25, both inflammatory and anti-inflammatory cytokines were elevated in the ASID cohort. **b)** Cytokineratio analysis of IL-2/IL-17, IL-12/IL-2, and IL-10/IFN-γ (left to right) at both time points provided greater resolution of inflammatory trends. Decreased ASID IL-17/IL-12 showed that despite increases in general inflammation, T cell proliferation did not occur. Additionally, IL-12/IL-2 was employed to investigate exhaustion, where increased antigen presentation (IL-12) did not stimulate T cell proliferation either. To rule out tolerogensis, IL-10/IFN-γ was employed, showing that ASID implantation did not evoke a response of that nature. **c**) Mice were implanted with either ASID (blue) or Mock Surgery (Black) on day 7 post-induction. By day 60 (after ASIDs had been completely resorbed), no ASID mice had relapsed, while two of three Mock Surgery mice displayed a fully severe recurrence of disease. (Statistical analysis was performed against the Mock group as a control. n = 3-4 biological replicates per group).

We contextualized cytokine trends across a number of mechanistically indicative ratios (**Fig. 7b**). IL-2/IL-17 was used as a measure of T cell proliferation proportional to overall EAE inflammation. Mock Surgery and Blank-treated mice showed increases in T cell proliferation at day 14 which was resolved by day 25, but ASID treatment appears to have restricted IL-2 at peak of disease. This trend is in concordance the exhaustive mechanism suggested by our other experiments. IL-12/IL-2 was used to interrogate the balance between APCs and T cells. Consistent with our earlier observations, ASID-treatment seems to cause an upward skew in this ratio, where APC levels are sustained while IL-2 is diminished. Finally, to investigate the observed increase of IL-10 from ASID splenocytes at day 25, IL-10/IFN-γ was employed to probe a potential tolerogenesis where anti-inflammatory IL-10 precludes pro-inflammatory IFN-γ production. Interestingly, despite these observed increases in IL-10, IFN-γ remains proportionally abundant such that this ratio fell well within the bounds of the controls.

Finally, we sought to discern whether ASID implantation would protect mice from the recurring paralysis that is characteristic of the PLP_139-151_-EAE model. In our early studies, ASIDs and Blank sponges alike were found to be completely resorbed by day 25. We wondered if a durable therapeutic effect could be realized as a result of ASID-induced cell exhaustion (**Fig. 7c**). During the initial phase of disease, only one of three ASID recipients displayed any EAE symptoms, and each of the Mock Surgery cohort exhibited a fully severe initial presentation. Remarkably, all ASID-implanted mice were protected from recurrent disease through day 60 while two of three Mock Surgery mice displayed robust relapses on days 37 and 38 post-induction. *Ex vivo* analyses revealed few differences between splenocytes in response to PLP rechallenge (**Supp. Fig. 8**).

## DISCUSSION

Previous works exploring antigen-specific immunotherapies for autoimmunity delivered autoantigen in vastly differing ways, including soluble, insoluble, and biomaterial compositions. ASIDs utilize covalent attachment of autoantigen onto a microporous scaffold to inhibit EAE *in vivo* when implanted subcutaneously after disease induction, but before the appearance of symptoms. Past work exploring immunologically-active therapeutic biomaterials has mostly explored co-delivery formulations [27]. To our knowledge, ASIDs represent a first demonstration of antigen-specific immune cell sequestration for therapeutic effect. Recently, the Serwold group disseminated an implantable biomaterial soaked in autoantigen for the purpose of homing and enriching antigen-specific populations in models of Type 1 Diabetes, but interestingly, no therapeutic effect was realized [35]. Our data also indicated antigen-specific cell infiltration into ASIDs (**Fig. 4, Fig. 5b**), but the therapeutic capacity we observed likely resulted from the specification of irreversible epitope conjugation to our constructs, which has been well described to enable antigen persistence and boost T cell stimulation [60]. In our example, such overstimulation proved itself a driver of effect through exhaustion and may explain our differential therapeutic success.

ASIDs elicited robust DC activation *ex vivo* (**Fig. 1e-g**). We initially hypothesized that this mechanism would solely facilitate the recruitment and sequestration of T and B effectors *in vivo*. When implanted, however, ASIDs spurred exhaustion and apoptosis within the constructs (**Fig. 3**). Evidence has emerged regarding the potential for “therapeutic exhaustion” by extrapolating a liability observed during chronic infection for possible benefit in autoimmunity [61]. In our studies, exhaustion was implicated in the prevention of disease. Spleen weights correlated highly with suppression of EAE, where engorgement was likely a consequence of ASID-induced exhaustion and return to SLOs (**Fig. 5b, Supp. Fig. 3** [14, 62]. The exhausted state of splenocytes in ASID-treated mice was corroborated by effector non-responsiveness to antigen rechallenge. Despite the presence of CD86+CD80+ active DCs (**Fig. 5d, e, and Fig. 6**), we observed an absence of IL-2 despite upregulation of other inflammatory markers such as IFN-γ (**Fig. 7a, b**). Interestingly, proliferation of resting (unchallenged) ABCs was diminished in the presence of PLP rechallenge (**Fig. 7c**). Abrogated proliferation was most severe in the antigen-specific population of this subset, suggesting that population changes may be driven by PLP-specific mechanisms. Autoimmunity was restricted to the periphery, exclusive of the CNS, as evident in the elevation of serum anti-PLP IgG, which was not observed in homogenized brain (**Fig. 3f**). In effect, the PLP-specific response was short-circuited and exhausted in ASID-treated mice, which translated to therapeutic success.

Remarkably, no ASID-implanted EAE mice developed significant disease (score >1) over 60 days. All three Mock Surgery-receiving counterparts exhibited severe primary EAE, and two of these mice exhibited robust flare-ups over this interval after day 25 (**Fig. 7c**). After the initial 25-day efficacy study, ASID and Blank constructs had been completely degraded, so this experiment was conducted to observe relapsed autoimmunity that is typical of the PLP_139-151_-EAE model. We originally expected ASID-treated mice to develop paralysis similarly to those given Mock Surgery by virtue of regained functionality after exhaustion, perhaps even at a higher rate or intensity of disease due to a delayed hypersensitivity reaction. In retrospect, initial disease prevention in ASID mice may have limited primary tissue inflammation and abrogated subsequent migration to the CNS.

Alternatively, ASID implantation could have induced differentially robust exhaustion similar to what is observed in chronic infection where clonal deletion or permanent anergy can ensue [18, 63–65]. In the EAE model employed here, immunity is generated against the PLP_139-151_ epitope, a small fraction of the larger PLP antigen, which is ultimately one of many constitutive building blocks of the myelin sheath. As a result, PLP_139-151_ density in the CNS is low and inaccessible relative to an ASID [66]. Conversely, PLP_139-151_ density in ASIDs is high and by design, easily accessible to extravasating immune cells. Antigen density is known to highly dictate costimulation and downstream immunity [66–69], where excessively high density can drive overstimulation and cell death [70–72]. Differences in these outcomes were evident between ASID-implanted mice and controls, especially in histological studies and bioassays of spongeocytes (**Fig. 4**).

In clinical practice, some of the most effective MS therapeutics such as natalizumab and fingolimod work by inhibiting immune cell migration and trafficking to the CNS [19, 20]. Such drugs have a steep trade-off, risking life-threatening consequences of nonspecific immunosuppression for therapeutic efficacy [21–23]. By demonstrating the potential to affect immune cell migration in an antigen-specific fashion, our work suggests autoimmunity may be short-circuited in potent, yet safer ways. Additional studies will determine if ASID performance can be extrapolated to a broader palette of autoantigens and if therapeutic efficacy can be achieved when intervening at different points in the disease. The immunological mechanism of ASID capture, deactivation, and release of exhausted autoimmune cells offers a new paradigm to short circuit immune cell migration to prevent relapsing autoimmune diseases.

## METHODS

### Materials

Microporous collagen sponges (4 mm and 21 mm, columnar pore architecture) were purchased from Advanced Biomatrix (San Diego, CA). 2,5-dioxopyrrolidin-1-yl 1-azido-3,6,9,12-tetraoxapentadecan-15-oate (azido-PEG4-NHS Ester) was purchased from Click Chemistry Tools (Scottsdale, AZ). Tris(3-hydroxypropyltriazolylmethyl)amine (THPTA), and sodium ascorbate (NaAsc) were purchased from Sigma-Aldrich (St. Louis, MO). Copper (II) sulfate pentahydrate (CuSO_4_·5H_2_O) was purchased from Acros Organics (Geel, Belgium). Alkyne-functionalized PLP with an *N*-terminal 4-pentynoic acid (homopropargyl, hp) modification, hpPLP_139-151_ (hp-HSLGKWLGHPDKF-OH) was purchased from Biomatik (Cambridge, ON, Canada). Unmodified PLP_139-151_ (NH2-HSLGKWLGHPDKF-OH) used for EAE induction, rechallenge assays, and anti-PLP IgG ELISA was purchased from PolyPeptide Laboratories (San Diego, CA). Incomplete Freund’s adjuvant (IFA) and killed *Mycobacterium tuberculosis* strain H37RA were purchased from Difco (Sparks, MD). Pertussis toxin was purchased from List Biological Laboratories (Campbell, CA). R-phycoerythrin (PE)/Cy7-conjugated anti-mouse CD3, PE-conjugated anti-mouse CD86, FITC-conjugated anti-mouse CD80, and respective isotype control antibodies were purchased from BioLegend (San Diego, CA). All other chemicals and reagents were analytical grade and used as received.

### Synthesis of ASIDs

Collagen sponges (4 mm or ¼ sections of 21 mm diameters) were modified by reacting in a 2 mg/ml solution of 2,5-dioxopyrrolidin-1-yl 1-azido-3,6,9,12-tetraoxapentadecan-15-oate (azido-PEG4-NHS ester) in 50 mM HEPES buffer at pH 8.3 for 4 hours at room temperature. Subsequently, the sponges were washed five times with deionized water. The washed sponges were then placed in an solution of N-(6-(diethylamino)-9-(2-(prop-2-yn-1-ylcarbamoyl)phenyl)-3H-xanthen-3-ylidene)-N-ethylethanaminium (Rhodamine-alkyne, 1.22 mM) or hpPLP_139-151_ (1.22 mM, 2 mg/mL) in pH 8.3 HEPES, followed by the addition of a premixed solution of tris-hydroxypropyltriazolylmethylamine (THPTA, 4.5 mM) and copper (II) sulfate pentahydrate (0.8 mM). Finally, sodium ascorbate (16 mM) was added to begin the reaction. The reaction was carried out overnight at room temperature. The sponges were then washed 5 times with deionized water and stored in 100% ethanol. The unmodified sponges were treated with the same procedure outlined above, with the omission of azido-PEG4-NHS ester. For *in vivo* studies, sponges were washed five times over the course of a day in 1X PBS and stored overnight at 4C in a solution of 1200 ng/mL of mouse GM-CSF.

### Characterizing ASIDs

PLP conjugation to collagen sponges was determined using a 20 minute Reverse-Phase HPLC method employing a 95/5 to 30/70 aqueous:organic gradient scheme on a C4 RP column. H_2_O + 0.05% trifluoroacetic acid was used as the aqueous phase, and acetonitrile + 0.05% trifluoroacetic acid was used for the organic. Functional characterization of ASIDs was completed by incubating PLP-conjugated and unconjugated sponges with 1:100 anti-PLP_139-151_ IgG positive mouse serum for 1 hour at room temperature. Sponges were washed, and bound anti-PLP_139-151_ IgG was detected with HRP-conjugated anti-mouse IgG.

### Induction of EAE and Therapeutic Study

EAE was induced as previously described [47, 59] in 4-6 week-old, female SJL/J mice from Envigo Laboratories. Mice were housed under specified, pathogen-free conditions at the University of Kansas with authorization approved under a protocol passed by the University’s Institutional Animal Care and Use Committee. EAE was induced by subcutaneous administration of 200 μg of unmodified PLP_139-151_ in 200 μL of Complete Freund’s Adjuvant (CFA) emulsion. The CFA mixture was produced from equal volumes of PBS and IFA containing killed Mycobacterium tuberculosis strain H37RA at a final concentration of 4 mg/mL. The immunization was administered as four, 50 μL injections above the shoulders and the flanks. Additionally, 200 ng of pertussis toxin was administered as an intraperitoneal injection on the same day of immunization (day 0) as well as day 2 post-induction. For therapeutic study, mice received a subcutaneous implantation surgery between the shoulder blades on Day 7. Mice were weighed each day of the study and monitored with clinical scores starting on day 7. Disease progression was assessed on a five-point clinical scale including: 0, no clinical evidence of disease; 1, tail weakness or limp tail; 2, paraparesis (weakness or incomplete paralysis of one or two hind limbs); 3, paraplegia (complete paralysis of two hind limbs); 4, paraplegia with forelimb weakness or paralysis; and 5, moribund.

### Detection of Anti-PLP IgG

PLP-specific IgG titers were assessed in an ELISA format as previously described [47, 48]. Briefly, 1μg/mL PLP was dissolved in a pH 9.5 solution of 50 mM NaHCO_3_. This coating buffer was seeded on Immulon 2HB 96-well plates and incubated overnight at 4°C. Plates were then blocked with 1% (w/v) bovine serum albumin (BSA). Serum and homogenized central nervous system tissue (hCNS) were then introduced and serially diluted 1:1 across seven concentrations starting at 1:100. After washing, HRP-conjugated anti-mouse IgG (BioLegend) was added at 0.1 μg per 100 μL. After a final washing regimen, 100 μL of TrueBlue substrate was added, and plates were covered and shaken at 250 rpm and room temperature for 15 minutes. Enzymatic conversion was stopped with 100 μL 2N H_2_SO_4_ and the plate was read at 450/540 nm (Spectramax M5, Molecular Devices). For analyzing titer, linear regions across sample titration readings were fitted with linear regressions and extrapolated to their 1X concentration for comparison across samples.

### Spleen Harvest and Splenocyte Isolation

Spleen harvest and splenocyte isolation was conducted as previously reported [57]. Briefly, spleens were passed through a wire mesh using the rubber stopper of a sterile 1 mL syringe in 1X PBS. The strained cellular extracts were centrifuged, and the cell pellet was resuspended in red blood cell lysis buffer. The cells were incubated on ice for 3.5 minutes to lyse splenic red blood cells. The lysis reaction was stopped by adding 10 mL RPMI 1640 media containing 10% FBS to the mixture before centrifuging. The remaining splenocyte pellets were resuspended in fresh media (RPMI 1640 media containing 10% FBS and 1% Penicillin-Streptomycin) and counted for further analysis and experimentation.

### Sponge Harvest and Spongeocyte Isolation

Cellular infiltrates to ASIDs and Blank Sponges were isolated by retrieving sponges from the subcutaneous space. In an Eppendorf tube, sponges were either chemically digested using 0.5 mg/mL Liberase TL (Sigma Aldrich, St. Louis, MO) in HBSS or mechanically disrupted with microscissors. The resulting freed cell suspension was then passed through a 70 μm strainer. The suspension was pelleted, replenished with 1 mL of RPMI 1640 media containing 10% FBS and 1% Penicillin-Streptomycin, and cells were counted for further processing and plating.

### Fluorescent Staining and Flow Cytometry

Splenocytes were collected from 24-well plates after 96 hours and stained with fluorescent antibodies according to manufacturer recommendations. Cells were washed with 1X PBS + 5% FBS + 0.1% sodium azide before being incubated for 20 minutes with Zombie Aqua viability stain (Biolegend) at room temperature. Following the incubation, fluorescent antibodies and isotype controls were added for 30 minutes on ice. For flow cytometry data collection, 100,000 cells per sample were targeted using a BD FACSFusion cytometer. Data were analyzed using FlowJo and GraphPad Prism software.

### Measurement of Cellular Metabolism

75 μM Resazurin (7-hydroxy-3H-phenoxazin-3-one 10-oxide) was incubated with splenocytes in a 96 well plate for 3 hours. Metabolic reductive capacity was observed by viewing changes in fluorescence at excitation 560, emission 590 (Spectramax M5, Molecular Devices). Background fluorescence was taken using RPMI media and subtracted out from splenocyte readings for analysis.

### Measurement of Cytokines

Following a 96-hour incubation, splenocytes in a 96-well culture plate were centrifuged. Supernatants were collected for cytokine analysis (GM-CSF, IFN- γ, IL-2, IL-21, IL-6, IL-10, IL-17, IL-23, TNF-α). Marker levels were detected using a U-Plex assay kit according to manufacturer instructions (Meso Scale Discovery). Each assay plate was read using the QuickPlex multiplex plate reader (Meso Scale Discovery).

### Histology

Resected sponges and tissues were cryoembedded with Tissue-Plus O.C.T. Compound (FisherSci) and sectioned into 20 μm on microscope slides in triplicate. Slides were fixed in 10% neutral buffered formalin for 10 minutes, and Masson’s trichrome stain (Sigma Aldrich) was conducted.

### Statistical Analysis

Statistical evaluation was performed using one-way analysis of variance (ANOVA), followed by Tukey and Sidak multiple comparison tests. Statistical significance for all analyses was set at p<0.05. All statistical analyses were performed using GraphPad Software (GraphPad Software Inc.).

## ACKNOWLEDGEMENTS

JDG was supported by the Madison and Lila Self Graduate Fellowship at the University of Kansas. JYS was supported by the Stella Fellowship of the Department of Pharmaceutical Chemistry at the University of Kansas. The authors would like to thank Dr. Steve Jacobson of the National Institutes of Health for his advisory role in study design, and for his continued interest in translating decoys for clinical applications. We would also like to thank Towne Walston, Sebastian Huayamares, and Michael Shao for their assistance in carrying out animal surgeries for this work, as well as Deanna Diaz for her help developing SAgAs as an antigen-specific B cell probe.

## DATA AVAILABILITY

The raw/processed data required to reproduce these findings cannot be shared at this time due to technical or time limitations.

