## Supplementary Information for "Antigen-Specific Immune Decoys Intercept and Exhaust Autoimmunity to Prevent Disease"

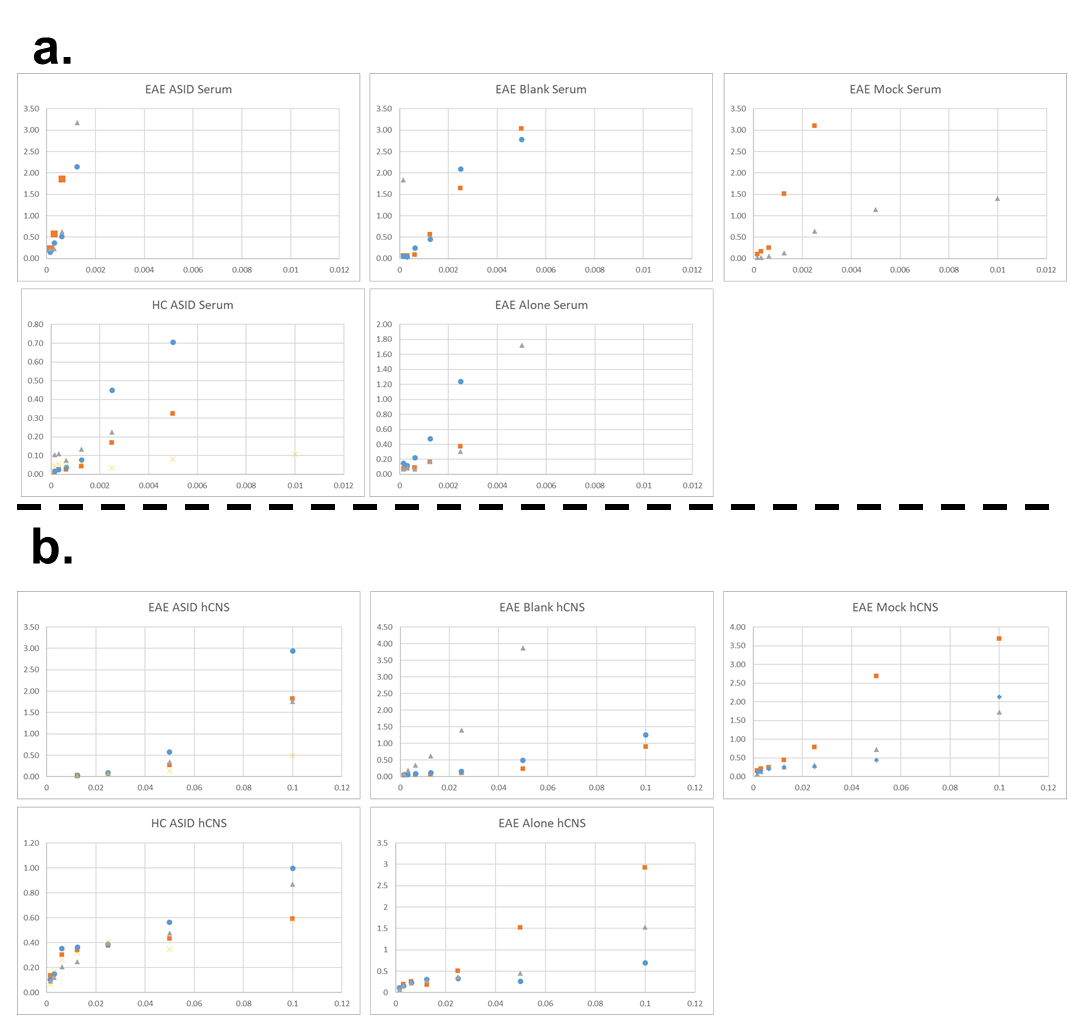

Supplementary Figure 1. Serum and hCNS autoantibody titers were detected by anti-PLP IgG ELISA. To compare relative titers, absorbance values were collected across seven dilutions, and the slopes of linear regions were calculated (r > 0.8) and compared. Both serum (**a**) and hCNS (**b**) were analyzed.

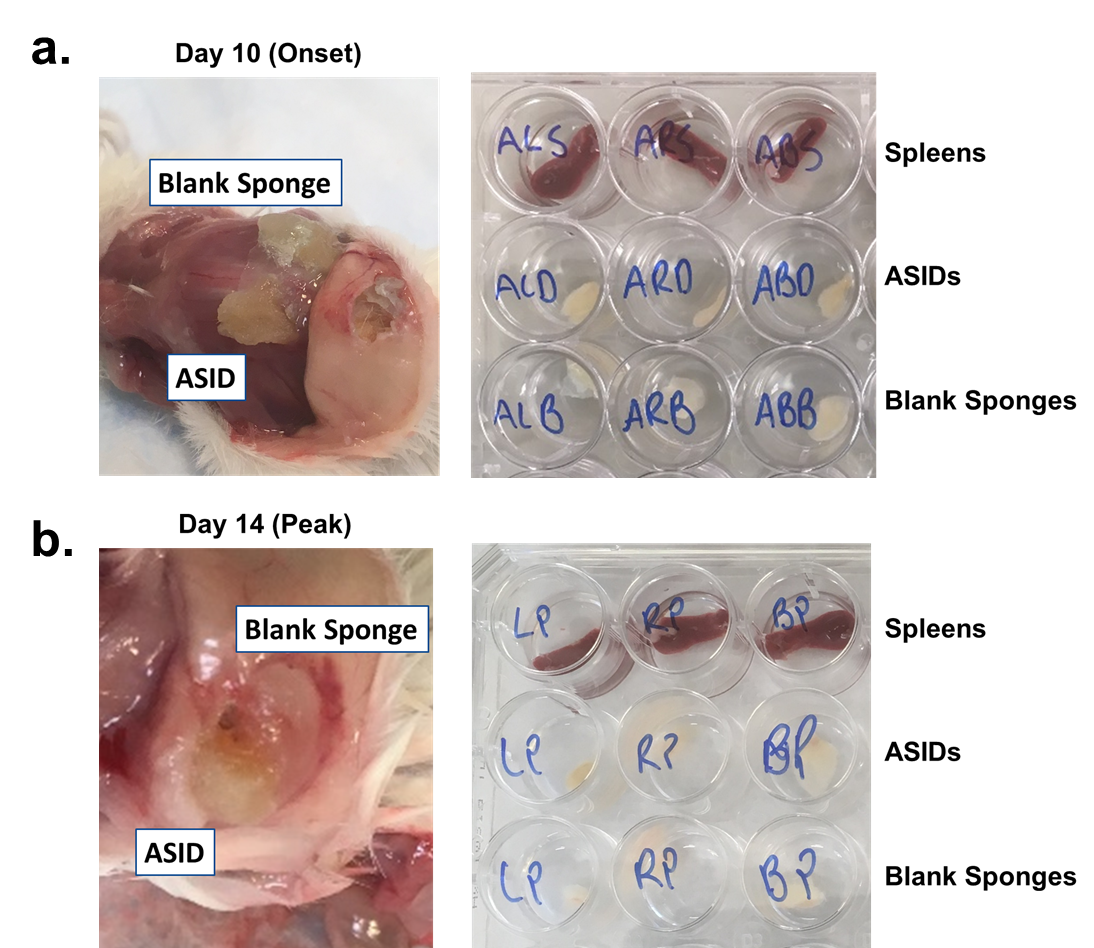

Supplementary Figure 2. In the same animal implant experiment, we observed engorged spleens at a time when secondary lymphoid organs are typically small. **a)** Day 10 and **b)** Day 14 sponges and spleens are pictured.

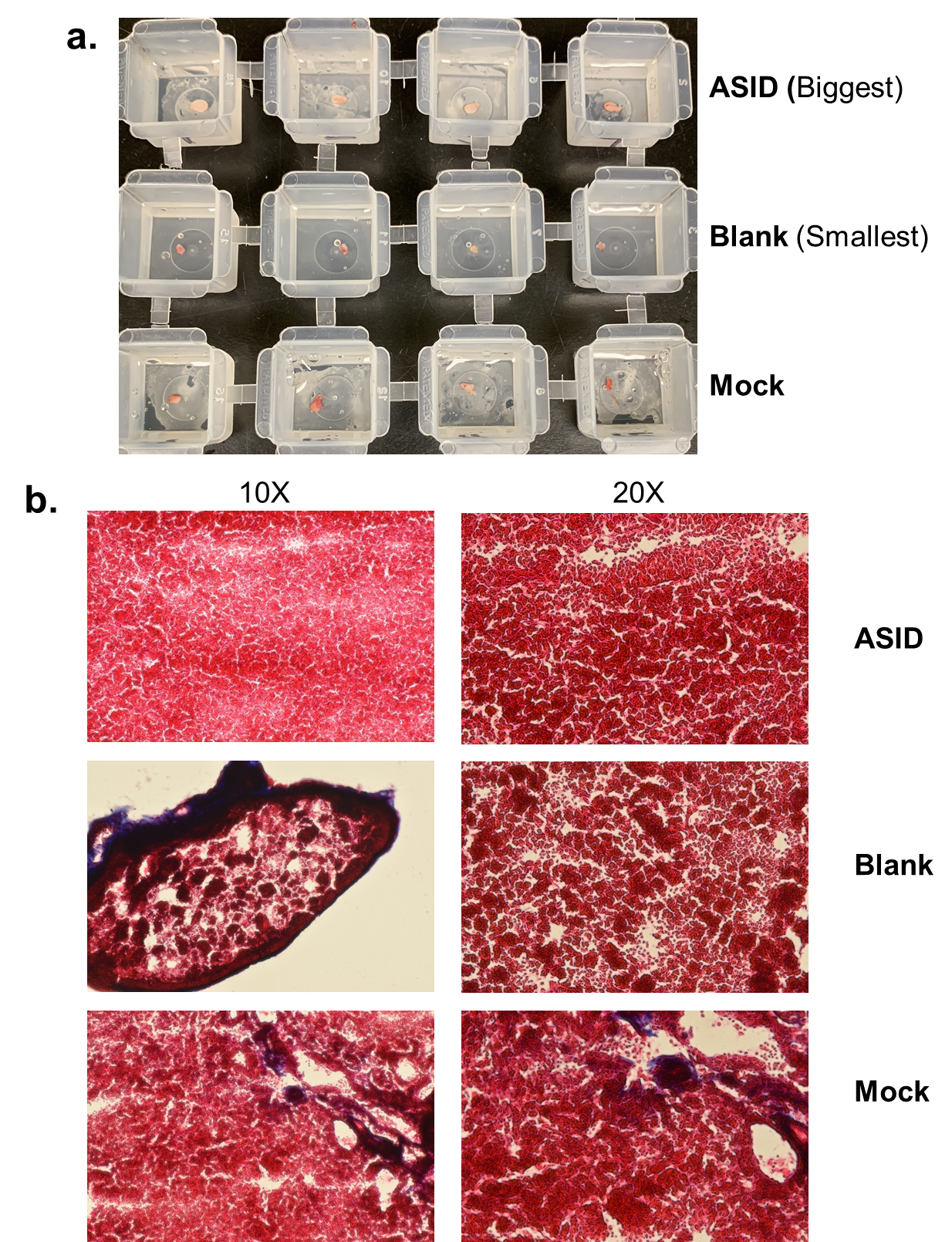

Supplementary Figure 3. EAE mice were implanted with either ASID, Blank Sponge, or Mock Surgery on Day 7 post induction. At day 14 (peak of disease), **a)** ASID popliteal lymph nodes were visibly swollen over Blank and Mock lymph nodes. **b)** Trichrome staining revealed a higher cell density in engorged ASID lymph nodes.

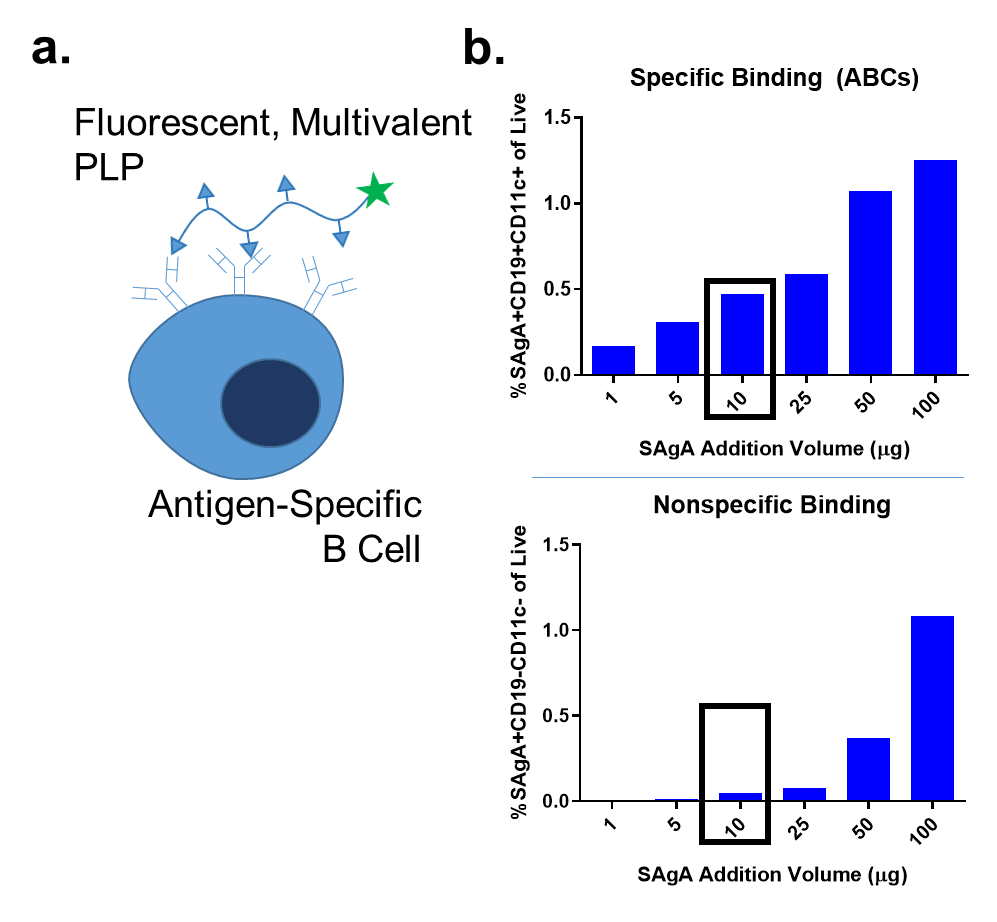

Supplementary Figure 4. Soluble Antigen Arrays (SAgAs) consist of autoantigen epitopes conjugated to a 16 kDa hyaluronic acid backbone. **a)** By incubating FITC-labeled SAgAs with EAE splenocytes, antigen-specific B cells were tagged through avid binding of the BCR. **b)** To determine a labeling method for fluorescent SAgAs, we titrated the amount of SAgA incubated per million splenocytes. Via flow cytometry, we measured antigen specific populations (ABCs, CD19+CD11c+) and nonspecific populations (CD19-CD11c-). 10 µg of SAgA per million splenocytes was selected as an appropriate labeling concentration, as it maximized specific binding while limiting nonspecific interactions.

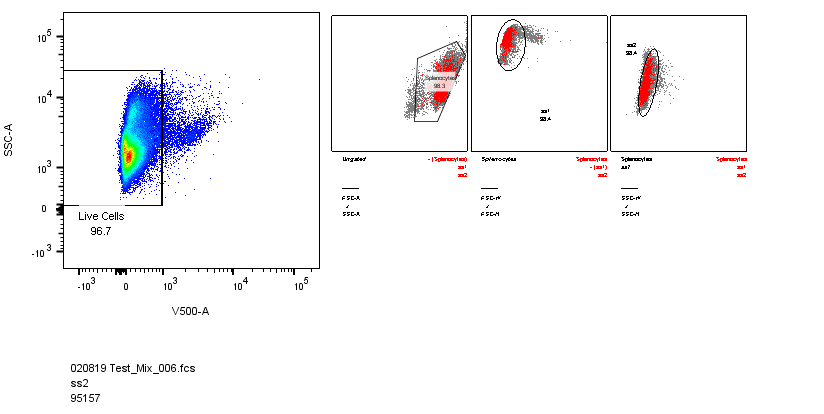

Supplementary Figure 5. Gating for flow cytometry live cell populations.

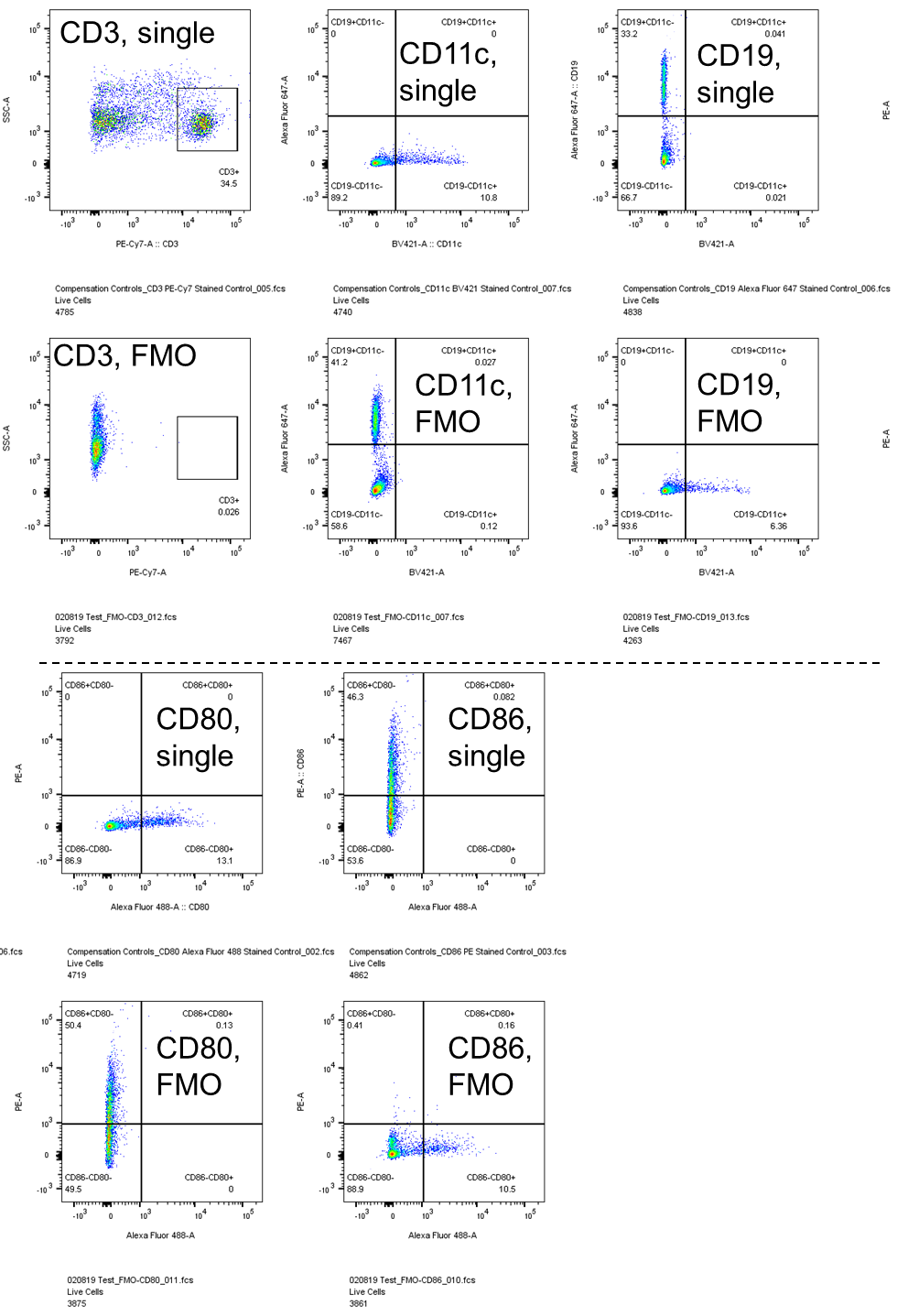

Supplementary Figure 6.. Gating for phenotypical populations. Shown are single stain controls (top) and FMO controls (bottom).

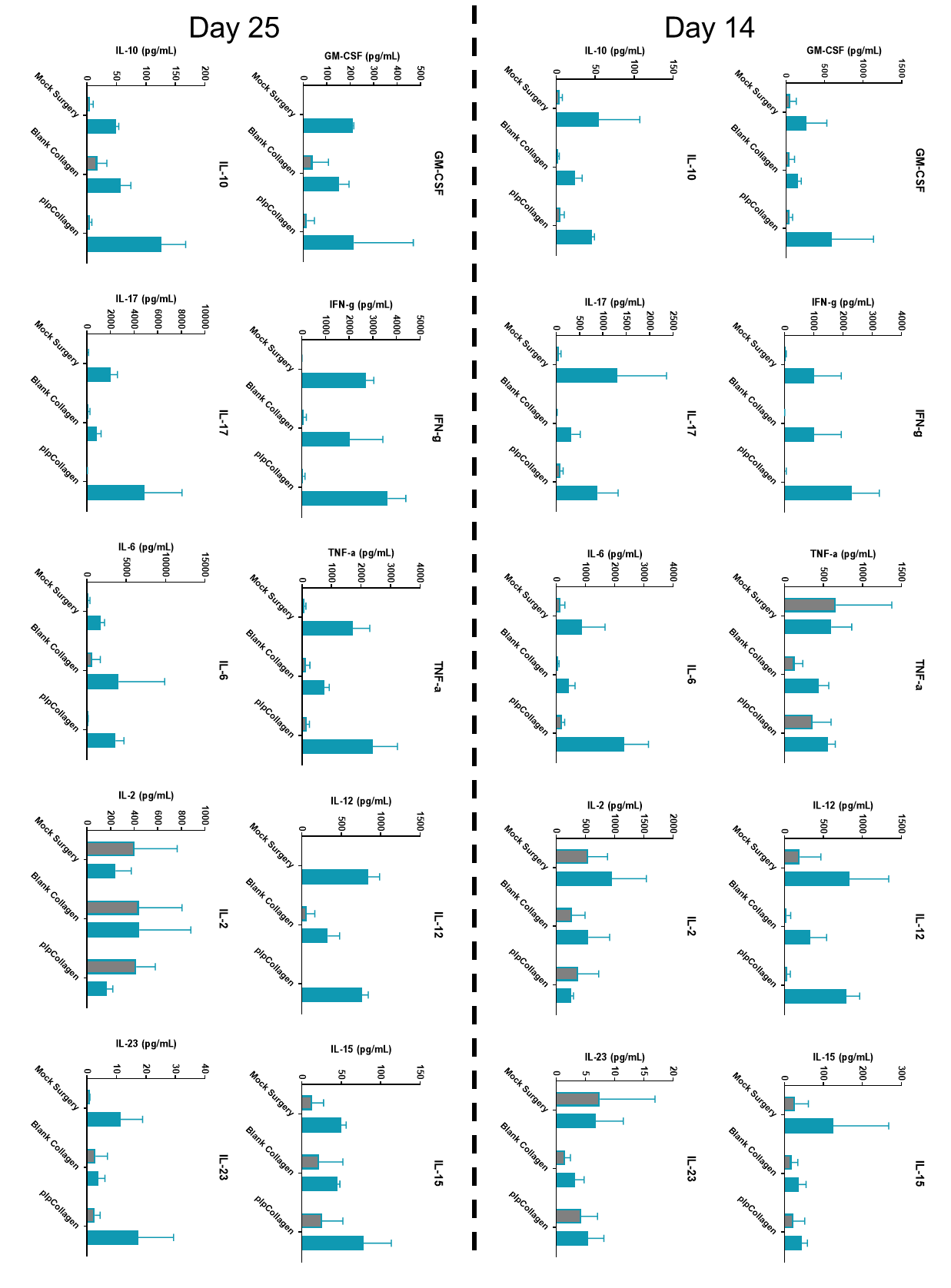

Supplementary Figure 7. Cytokine Analyses are presented as absolute pg/mL values to supplement heatmap results presented in Figure 7.

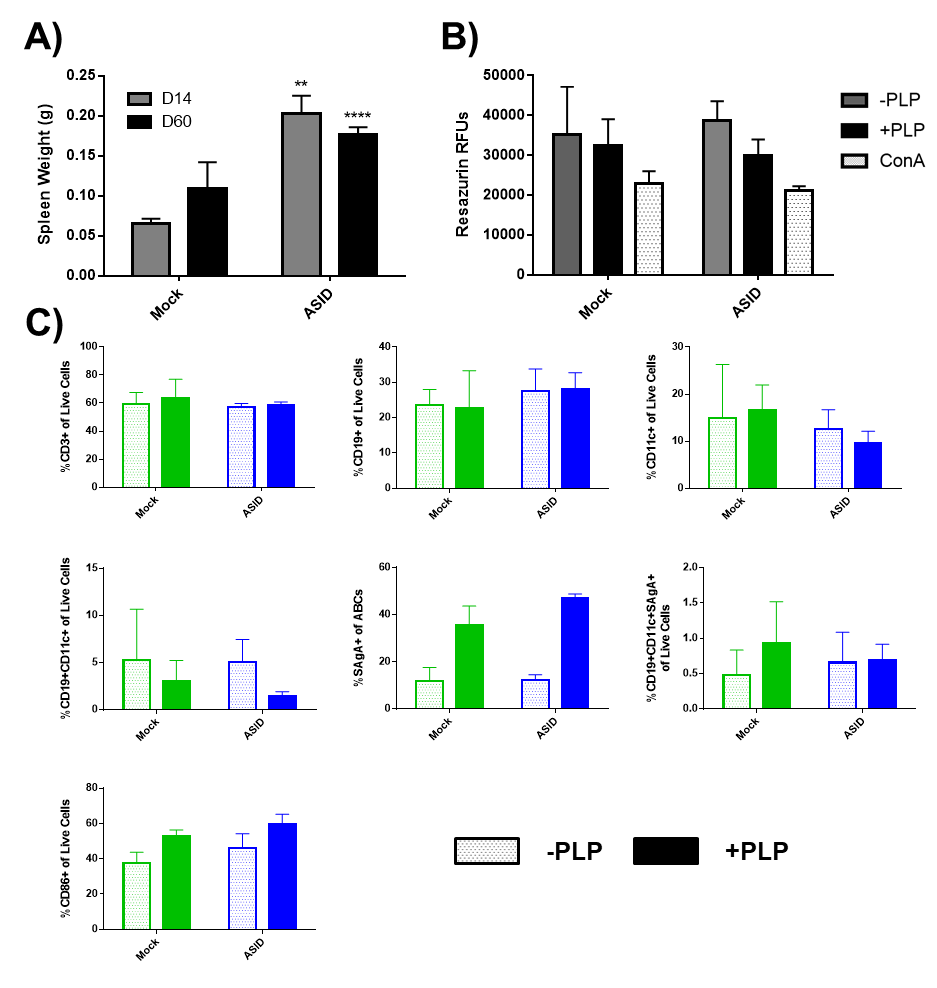

Supplementary Figure 8. PLP rechallenge response among harvested D60 splenocytes. **A)** Spleen weights were taken upon harvest and compared to D14 observations. **B)** Resazurin cell metabolism was measured after 96 hours of incubation with –PLP, +PLP, or ConA. **C)** Splenocytes were phenotyped after 96 hours of vehicle (-PLP, open bars) or 25 µM PLP rechallenged (+PLP, solid bars) for CD3, CD19, and CD11c (top row, left to right) as well as antigen-specific B cell populations (middle row), and costimulation (CD86, bottom row). n = 3 biological replicates per group, *p < 0.05, **p < 0.01, ***p < 0.001, ****p<0.0001).
